# A shift in the host web occupancy of dew-drop spiders associated with genetic divergence in the Southwest Pacific

**DOI:** 10.1101/2023.11.13.566795

**Authors:** Noraya U. Elias, Mae A. Responte, Cheng-Yu Wu, Yi-Fan Chiu, Po Peng, Hauchuan Liao, Rafe M. Brown, Yong-Chao Su

**Affiliations:** Department of Biomedical Science and Environmental Biology, College of Life Science, Kaohsiung Medical University, Kaohsiung, Taiwan; Mindanao State University-Malabang Community High School, Malabang, Lanao del Sur, Philippines; Graduate Institute of Medicine, Kaohsiung Medical University, Kaohsiung, Taiwan; Department of Biological Sciences and Environmental Studies, College of Science and Mathematics, University of the Philippines Mindanao, Davao City, Philippines; Systematic Zoology Laboratory, Department of Biological Sciences, Tokyo Metropolitan University, Tokyo, Japan; Biodiversity Institute, Department of Ecology and Evolutionary Biology, University of Kansas, Lawrence, Kansas, USA

**Keywords:** Geogenomic framework, host adaptation, Kerama gap, kleptoparasite, RAD-seq genotyping, long distance dispersal

## Abstract

**Aim:** We assessed the population genetic structure of the kleptoparasitic spider *Argyrodes bonadea* across the Southwestern Pacific islands. Our focus is on assessing the impact of overseas distances and, in particular, the Kerama gap, as potential drivers of genetic differentiation. We found that the spider kleptoparasite’s switch to a specific host species is associated with significant genetic variation at fine scales, whereas the same species adoption of a generalist host strategy has likely facilitated its broad dispersal, colonization, and recent range expansion across the southwestern Pacific, and is associated with a lack of geographically– structured genetic variation in these latter, subsequently-colonized landmasses.

**Location:** Southwestern Pacific Islands

**Taxon:** *Argyrodes bonadea*

**Methods:** We used mitochondrial Cytochrome Oxidase 1 (CO1) gene sequences, and Restriction Site-associated DNA Sequencing (RAD-seq) for our analyses.

**Results:** Two strongly supported lineages, an Amami-Okinawa Lineage (AOL) and an Austral-Asia Lineage (AAL) correspond to two separate clades, roughly divided by the Kerama Gap, in phylogenetic trees estimated here. However, species delimitation led to the interpretation of only a single species present. The AOL exhibits complex, geographically-structured host web spider species specificity, wherein the Amami population utilizes *Cyrtophora*, but AOL samples in Okinawa associates exclusively with *Nephila*—and yet all broadly distributed AAL populations show no evidence of host web spider species specificity.

**Main conclusion:** The population boundary between AOL and AAL likely results from local adaptation to novel hosts—instead of isolation by the Kerama Gap—following long-distance dispersal and range expansion. Our results suggest kleptoparasitic spiders have the capacity to overcome permanent deep-sea barriers and colonize distant landmasses. Whereas peripheral populations (AOL) demonstrate the capacity for specialization to a single host, which may have contributed to genetic differentiation and isolation, the broadly-distributed AAL persists and has successfully expanded its geographical range as a host generalist, which may contribute to ongoing gene flow inferred in this study.

## INTRODUCTION

Since the early 1980s, vicariance theory emerged as the dominant paradigm for testing hypotheses in biogeography (Nelson & Platnick, 1980), making geographic isolation the widely-adopted null hypothesis influencing distribution patterns and differentiation processes in many taxa (Wiley, 1988; Sampson et al., 1998; Upchurch, 2008; Baker, Boyer, & Giribet, 2020; Tolley, Tilbury, & Burger, 2022). Adherence to this polarized argument (e.g., Heads, 2005) may hinder progress toward considering and understanding the importance of many other factors (De Queiroz, 2015) such as long-distance dispersal (Welt & Raxworthy, 2022) and the role of ecological adaptation (Zhou, Tran, Kieft, & Anantharaman, 2020), which often contribute to unique floral and faunal assemblages of celebrated island communities (Wallace, 1869; Wagner & Funk, 1995; Gillespie, 2002; Lomolino, Riddle, & Whittaker, 2017). In East Asia, the Ryukyu Island Chain had been studied extensively, as a fertile region to test the relative influence and impact of geohistorical events, deep-sea barriers, and sea currents in shaping this archipelago’s highly-endemic, insular biota (Kawamura, Chang, & Kawamura, 2016; Yang, Komaki, Brown, & Lin, 2018; Komaki, 2021). However, ecological factors such as host-specificity, adaptation to local environments, and interspecific competition had been rarely considered (but see Sato et al., 2021). Furthermore, the limited available studies have primarily focused on Taiwan and the Ryukyus with occasional attempts to establish connections with continental biota (Yang et al., 2018; Chen et al., 2021). Thus, exploring distant regions beyond nearby islands, such as the Philippines, have the potential to uncover hidden species distribution patterns and provide new opportunities for testing genetic variation using genomic data.

We characterized genomic variation in a model study system, leveraging the unique host-web parasitism modes of kleptoparasitic spiders of Taiwan, the Ryukyu Island Chain, the Philippines, and Australia. The unusual natural history of our focal taxon allows us to test the prevalence, and evaluate the relative importance of historical biogeographic processes versus ecological factors, which may act variably on lineage diversification in the insular faunas of East Asian archipelagos using a geogenomic approach (Dawson, Ribas, Dolby, & Fritz, 2022). Geographic isolation has long been considered to be the most prevalent factor in the process of speciation (Darwin,1859; Mayr, 1963; Nelson & Platnick, 1980; Sobel, Chen, Watt, & Schemske, 2010). Ancient continental breakup and widespread global vicariance, widely adopted to explain the division of faunas, has also been invoked as likely leading to blockage of dispersal, cessation of gene flow among populations, and the predominant explanation for disjunct distributions (Nelson & Platnick, 1980; Gillespie et al., 2012; Lomolino et al., 2017). Without dispersal and gene flow, isolated populations tend to diverge, often leading to speciation (Mayr, 1947; Volk, Konvalina, Floeter, Ferreira, & Hoffman, 2021). The Ryukyu Island Chain is a biologically unique continental arc, located between Japan to the north, and Taiwan to the south. This island chain is composed of continental fragments that lie on the western rim of the Pacific Ocean (Chiang & Schaal, 2006), and are geographically divided into northern, central, and southern regions by two deep-sea barriers: The Tokara and Kerama Gaps, respectively (Kizaki, 1978; Ota, 1998). These formidable, permanent, deep-water barriers kept each region isolated from each other since the late Miocene to the early Pleistocene (7.0–1.7 MYA, Ota, 1998; Osozawa et al., 2012). Even during the Pleistocene glacial cycles with lowered sea levels c. 85– 130 m (Peltier, & Fairbanks, 2006; Zhai, Comes, Nakamura, Yan, & Qiu, 2012), land bridges likely did not form across the Tokara and Kerama Gaps due to their depth exceeding 1000 m. (Kimura, 2002; Inoue et al., 2020). The formation of permanent marine barriers to dispersal by terrestrial animals, and the repeated cycles of connection and disconnection of several smaller islands (separated by channel depths of ≤ 120–180 m) within each region during the Last Glacial Maxima (LGM) has been interpreted as having contributed strongly to the assembly and formation of the Ryukyu Archipelago’s endemic flora and fauna (Nakamura et al., 2015; Chiu et al., 2017; Shen, Chang, & Ota, 2022). During the Pleistocene, the southern portion of the Ryukyus was repeatedly linked to Taiwan via ephemeral land bridges (Kimura, 2000). Extending to the southern limit of Taiwan lies a small chain of volcanic islands, extending between Taiwan and the Philippines, with no evidence of connection during the Pleistocene (Batanes–Babuyan island groups; Heaney, 1985; Brown et al., 2013). Thus, dispersal between the Philippines and Taiwan, and colonization of fauna in both directions is often attributed to the classic “stepping stone” (Ota, & Huang, 2000; Oliveros, Ota, Crombie, & Brown, 2011) or island-hopping mode of successive dispersal events (Esselstyn & Oliveros, 2010; Layos, Geromo, Espina, & Nishibori, 2022).

The geogenomic approach allows for critical evaluation of a hypothesis testing framework, by way of evaluating alternative predictions using genomic DNA sampled from across the genome (Dolby et al., 2022). Recent improvements in high-throughput and reduced representation genomic techniques such as Restriction Site-associated DNA Sequencing (RAD-seq; Andolfatto et al., 2011) are enabling a cost-effective use of genomic fragments, by targeting single nucleotide polymorphism (SNP) markers for inferring population genetic parameters (Andrews, Good, Miller, Luikart, & Hohenlohe, 2016). These markers are the most common sources of genetic variation among members of the same species and currently are the focus of genome-wide research (Uffelmann et al., 2021). Cross-genome information represented in SNPs data is powerful for species delimitation (Georges et al., 2018; Arrigoni et al., 2020; Ivanov, Marusik, Pétillon, & Mutanen, 2021), distribution patterns (Colli et al., 2018), molecular ecology, and geographically-explicit landscape characterizations of organismal dispersal conduits and corridors (Dömel et al., 2020)—especially for characterizing these patterns intraspecifically or among closely-related species. However, RAD-seq proves less efficient in evaluating hypotheses related to adaptation and selection, given its susceptibility to miss loci under selection in investigations of local adaptation (Lowry et al., 2017).

*Argyrodes* spiders are kleptoparasites that evolved from araneophagic ancestors to adopt group-living kleptoparasitic lifestyles (Su & Smith, 2014). Kleptoparasites inhabit the host spider webs and rely exclusively on these webs for obtaining their food. Their feeding niches and distributions are constrained by geographic boundaries and host availability. However, the relative importance of these variables in population persistence, dispersal, colonization, and evolutionary diversification remains unexplored. Here, we assess the population genetic structure of the kleptoparasitic spider *Argyrodes bonadea* Karsch, 1881(Levi & Levi, 1962), which is co-distributed with its large orb–weaving spiders across Australia, the Philippines, Taiwan, and the Ryukyu Island Chain. The host spider web is vital to the life history of *A. bonadea*: within the web, females deposit their egg-sacs to ensure access to food for their spiderlings, which persist in the web until adulthood. Despite the unusual natural history and reproductive mode, the genetic structure of this kleptoparasite has not been fully studied. However, it is possible that it exhibits a similar pattern of limited geographic genetic variation if geographic barriers do not effectively prevent or restrict gene flow, as observed in its unrelated host species (e.g., *Nephila pilipes,* Su, Chang, Lee, & Tso, 2007). To investigate this possibility, we genetically assessed populations of *A. bonadea* across the long Ryukyu island chain and evaluated two general predictions. We assumed allopatric differentiation of kleptoparasite lineages in which more pronounced differentiation would be expected in the most isolated populations across the islands of Ryukyu, Taiwan, the Philippines, and Australia. Under this island barrier-driven differentiation expectation, divergence times should be similar to the dated geological events which underly the causal mechanisms and history relating to how and when landmasses were isolated (e.g., >7.0 MYA for populations in the Ryukyus, when they started becoming isolated from the continent (Kimura, 2002). In contrast, if marine barriers have been ineffective as geographic isolating factors, genomic variation detected here—and underlying divergence—may alternatively be generated through non–vicariant mechanisms, such as dispersal following adaptive processes; thus, kleptoparasite lineages may have diverged later, when they specialized subsequently to their respective host species. We performed phylogenomic inference–assisted species-delimitation analyses, to gain insight into the number of likely species–level (differentiated) lineages represented by the focal kleptoparasites, to provide context for our interpretations, regarding “specialist” versus “generalist” kleptoparasite spider population natural histories. Finally, we utilize our geogenomic framework, a novel empirical genomic dataset, and new kleptoparasite-host association data empirically gathered in the field, to demonstrate for the first time that a switch to a specific host species by a kleptoparasitic spider leads to significant genetic variation at fine scales. However, our study species’ adoption of a generalist host strategy appears to have facilitated broad, long-distance dispersal, colonization, and recent range expansion, resulting in a lack of geographically-structured genetic variation in subsequently colonized regions.

## MATERIALS AND METHODS

### Taxon collection

We collected samples from the Philippines, Australia, Ryukyu Island Chain, and Taiwan during 2007 to 2020 (Fig. 1a). The areas covered by our sample collection including northern Palawan and Mindanao (Surigao Province) islands, Philippines; Queensland, Australia; the Ryukyu Island Chain that consists of Yaeyama, Miyako (includes Irabu and Ikema), Okinawa, and Amami; Taitung, Taiwan. During fieldwork targeting *A. bonadea*, we recorded individual host identities based on co-inhabiting spiders found in the same web and functional association with each *A. bonadea* individual. Data without a cohabiting host or cases of one *A. bonadea* + two or more host genera were excluded from our analysis. Web-building hosts included different species of the genera *Argiope, Cyclosa, Cyrtophora*, *Leucauge, Neoscona, Nephila,* and *Parasteatoda*. We collected a total of 112 *A. bonadea* samples, all with complete host association data (Appendix S1); seven cases of ambiguous or uncertain host association were discarded. All the fresh specimens were preserved in 95% ethanol and stored at −30°C until further use. Sample storage, collection of morphological and molecular data was carried out in Ecology and Evolutionary Genomics (EEG) Laboratory at Kaohsiung Medical University, Kaohsiung, Taiwan.

**Figure 1.**
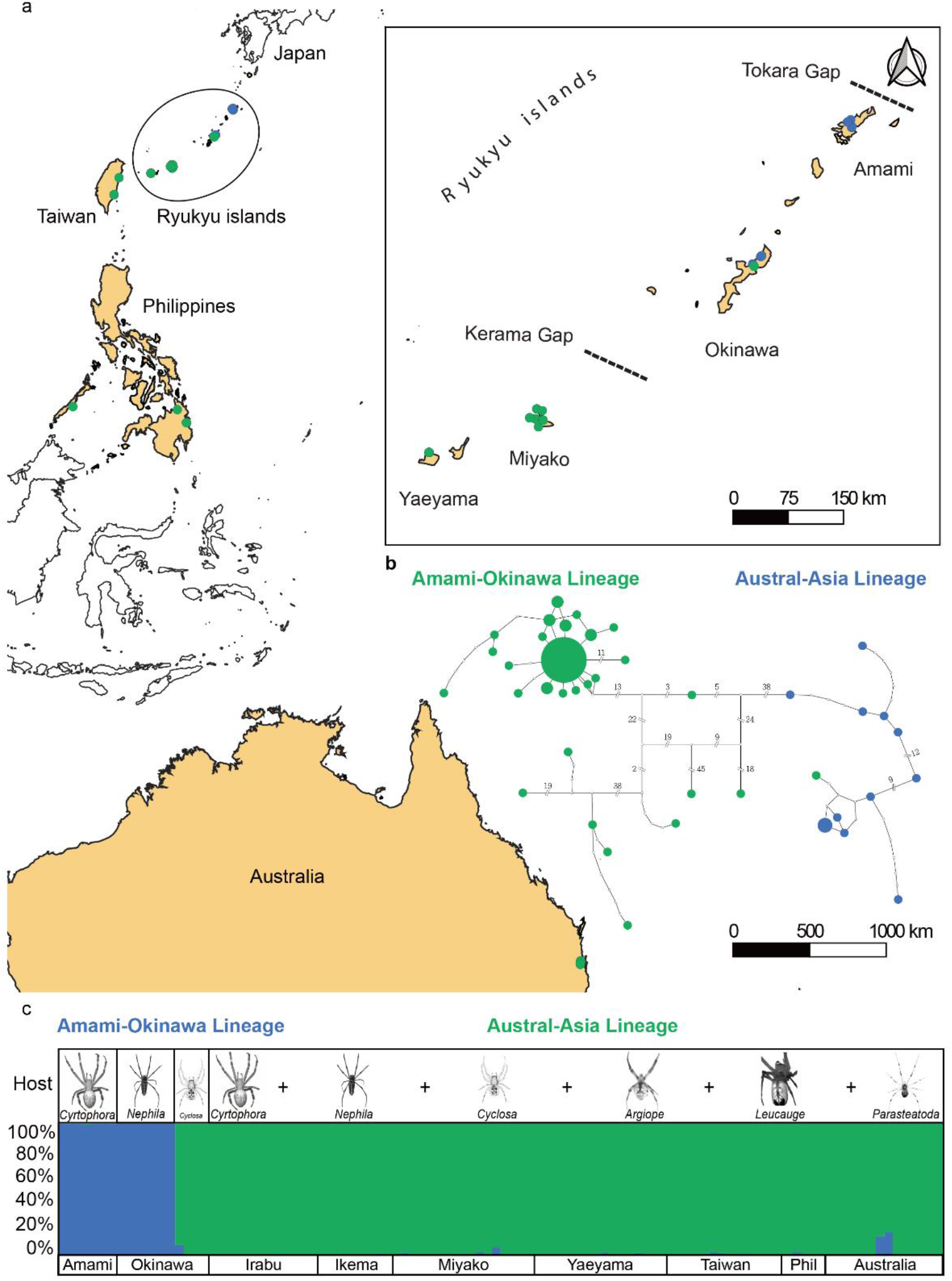
(a) Collection sites and geographical range of the primary study species. (b) Haplotype network shows geological relationships of different populations using CO 1 data, where in mutational steps correspond to substantial geographical distances, presumably reached via dispersal. (c) Population genetic STRUCTURE analyses of single-locus single nucleotide polymorphism (SNPs) data revealed two lineages: the Amami–Okinawa Lineage (AOL) showing host-specificity and Austral–Asia Lineage (AAL) utilizing a broader variety of different hosts.

### Morphological variation

We assessed morphological variation among samples from different islands focusing on measurement of continuously varying morphometric body dimensions and characters. Character images were captured using a Nikon camera D5600 mounted on a Leica stereomicroscope and measurements were recorded from digital media, using GIMP 2.10 software (GIMP Team, Creative Commons Attribution-ShareAlike 4.0 International License). Individual measurements of 27 and 26 characters in males and females, respectively, were recorded using character definitions of Kim & Kim (2007). Following Responte et al. (2021), data from different islands, and for males versus females, were analyzed separately to avoid potential confounding variation due to size and shape sexual dimorphism. We performed the Principal Component Analysis (PCA) using the “prcomp” function in R 3.6.1 (R Core Development Team, 2019) and calculated the Eigenvalues that correspond to the amount of morphological variation by each principal component. The first two components with the highest Eigenvalues where extracted and used as the final result in the PCA plot which were visualized using the R package *ggfortify* (Horikoshi & Tang, 2018).

### Genomic DNA extraction, PCR amplification, and sequence alignment

DNA samples were extracted from the legs of adult specimens and the prosoma of sub-adult specimens. We followed the procedural steps of Maxwell® RSC Blood DNA Kit AS1400. We used 300 μl of Tissue Lysis Buffer (TLA) instead of the lysis buffer provided in the extraction kit and 30 μ*l* proteinase K (PK) then incubated at 56°C overnight for homogenization. Genomic DNA was purified using the Maxwell® RSC Instrument following the standard manufacturer’s instructions then eluted in 50 μl buffer and quantified using Quantus Fluorometer (E6150, Promega Corp., USA). The purified genomic DNA was stored at −30°C until subsequent Polymerase Chain Reaction (PCR) amplification and genomic library preparation. We followed the same procedure for amplifying the Cytochrome Oxidase 1 (CO1) fragment as in Responte et al. (2021). For the RAD-seq library preparation and sequence assembly, we used the methods described by Andolfatto, Davison, Erezyilmaz, Hu, Mast, Sunayama-Morita, & Stern (2011) and Catchen, Hohenlohe, Bassham, Amores, & Cresko (2013). The purified final products of the targeted 375 base pairs were sequenced using Illumina NovaSeq 6000 sequencing with 2 × 150 bp kit. We used Stacks v2.0 (Catchen et al., 2013) to assemble the raw RAD-seq sequences using default parameters and loci were filtered out at 50% occupancy rate. The details are described in Appendix S2.

### Population structure and haplotype network

We employed the STRUCTURE analysis (Pritchard, Stephens, & Donnelly, 2000), a Bayesian clustering approach using K-mean algorithm, to initially categorize all individuals from a given sample into an ancestral population membership based on the assumption that genetic similarity among individuals is structured by the existence of subgroups and geographic isolation (Novembre, 2016). We ran the Cluster Markov Packager Across K (CLUMPAK) using the main pipeline to find the best K modes with 10 independent runs for each value of K ranging from 1 to 10. The optimal K value was calculated according to Evanno, Regnaut, & Goudet (2005) and the final output was visualized in Excel.

To compare the results of the SNPs in STRUCTURE with simple measures of genetic variation using our mtDNA CO1 marker, we calculated nucleotide and haplotype diversity using DnaSp 6.12.03 (Rozas et al., 2017). To identify haplotypes and their genealogical relationships, we generated haplotype networks in TCS 1.21 (Clement, Posada, & Crandall, 2000) and visualized them using tcsBU (Múrias dos Santos, Cabezas, Tavares, Xavier, & Branco, 2016).

### Population genetic clustering

We also performed the Discriminant Analysis of Principal Components (DAPC; Jombart, Devillard, & Balloux, 2010) using the K-mean algorithm and the Bayesian Information Criterion (BIC) to identify the ideal number of clusters with ADEGENET package (Jombart, 2008) in R studio 4.1.0. The Variation Autoencoder (VAE; Derkarabetian, Castillo, Koo, Ovchinnikov, & Hedin, 2019) was also used to further visualize sample clustering. This unsupervised machine learning method is derived from Bayesian probability theory and recently has been shown to be effective for species delimitation (Derkarabetian et al., 2019; Newton, Starrett, Hendrixson, Derkarabetian, & Bond, 2020). The pipeline consists of an encoder that takes in the one-hot file SNPs as unlabeled input data, and performs a training phase in which the system learns patterns in the data. A subsequent decoder stage then reconstructs and clusters the output, based on major features learned from structure in the original data. The final output is a two-dimensional graphical representation of the clustered SNPs, modeled according to the structure in the original data.

### Phylogenetic analyses

#### BEAST Tree for CO1 and RAD-seq

We employed *A. fissifrons*, *A. lanyuensis*, and *A. miniaceus* as outgroup species, demonstrating their close relation to *A. bonadea* through molecular phylogenetic analysis (Su and Smith, 2014) and RAD-seq data with known distribution (Responte et al., 2021; Platnick, 2023). The best fit model was calculated based on Akaike Information Criterion (AIC) with the use of the jModelTest2 v. 2.1.1 (Darriba, Taboada, Doallo, & Posada, 2012) in CIPRES (Miller, Pfeiffer, & Schwartz, 2010). The general time reversible (GTR) model was used as the most suitable model for both the CO1 and RAD-seq data. The XML file was generated in BEAUTi 1.10.4 (Drummond, Suchard, Xie, & Rambaut, 2012) and Yule process speciation was used as our tree prior then uncorrelated relaxed-clock was applied with an assumption that each branch of the phylogenetic tree evolved independently (Drummond et al., 2012). For the ucld.mean we applied the lognormal prior distribution with the initial value (0.1), mean value (0.0112), and a standard deviation value (0.01) for the CO1 data (following Su & Smith, 2014), but for the RAD-seq data we used the uniform flat prior. Markov Chain Monte Carlo (MCMC) chain length was set at 5 × 10^8^ (CO1 data) and 1 × 10^9^ (RAD-seq data) with sampled trees saved every 1 × 10^3^. The generated XML file was run in BEAST v.1.10.4 (Drummond et al., 2012) with three replicates. The log files were combined, then the trace file results were visualized in Tracer v.1.7.1(Rambaut, Drummond, Xie, Baele, & Suchard, 2018) to determine the effective sample size (ESS > 200) and discard the first 10% of the sampled trees as burn-in. We generated the maximum clade credibility (MCC) tree in TreeAnnotator v.1.8.4 (Rambaut & Drummond, 2010), and visualized it using FigTree v.1.4.3 (Rambaut, 2014).

#### IQ-TREE 2, Bootstrap Support, Gene Concordance Factor (gCF), and Site Concordance Factor (sCF)

To quantify the gene concordance factor (gCF) and site concordance factor (sCF), we constructed the maximum likelihood (ML) reference tree in IQ-TREE 2 (Minh et al., 2020). The gene concordance factor (gCF) measures how many percentages of the gene trees agree with the reference tree for each specific branch while the sCF calculates the percentage of agreement of individual alignment sites with the reference tree in that branch (Minh et al., 2020). These measures of branch support allow us to consider conflicting topologies with strong support from the same datasets, as seen in studies across various organisms (Cloutier et al., 2019; Kulkarni et al., 2021). This strongly suggests that while bootstrap and Bayesian posterior support are crucial in phylogenetic analyses, they may overlook inherent variation within genomic data, where each gene and site can possess a distinct evolutionary history. We used the assembled loci in Phylip format to reconstruct our reference species tree with 1 × 10^3^ ultrafast bootstraps. We constructed gene trees using the -S command to compute the individual locus. We computed the gCF and sCF separately using the reference species tree and the concatenated gene trees, then combined the two output files in a single run. The final output was visualized in FigTree.

#### Divergence-time estimation with SNPs data

We employed a method for time calibration that combines a molecular clock model, which can be calibrated with fossil or biogeographical constraints with Bayesian species-tree inference of the program SNAPP (Bryant, Bouckaert, Felsenstein, Rosenberg, & RoyChoudhury, 2012). The single-locus SNP Phy format is used as the input file, and we follow the workflow proposed by Stange, Sánchez-Villagra, Salzburger, & Matschiner (2018) to generate the XML file. To specify the crown age constraints, a normal distribution was used with 20.10 MYA as mean and 0.01 as standard deviation based on the divergence of the outgroup and ingroup (Su and Smith, 2014). The outgroup clade was treated as a monophyly constraint with the application of “monophyletic, NA.” Under TreeAnnotator, we summarized an MCMC chain length of 1 × 10^5^ with a burn-in of 1 × 10^4^. We targeted the tree type as maximum clade credibility tree and selected the node heights by the mean.

#### Bayes Factor Delimitation (BFD)

We tested three species hypotheses models: model 1 (one species model, all *A. bonadea* lineages lumped as one species), model 2 (two species model, based on phylogenetic results, AOL and AAL considered as two different species), model 3 (three species model, Amami, Okinawa1, versus remaining island populations), model 4 (four species model, Amami, Okinawa1, Okinawa2, and remaining island populations) through Bayes Factor Delimitation (BFD, Kass & Raftery, 1995). This method gives an estimate of the marginal likelihood of each of the pre-defined species hypotheses models implemented in SNP and AFLP packages for phylogenetic analysis (SNAPP) (Leaché, Fujita, Minin, & Bouckaert, 2014) under BEAST2. The input data are single locus SNPs in nexus format, to generate an XML file in BEAUTi2, for each of the four species hypotheses models executed in BEAST2—with 48 path sampling steps, an alpha of 0.3 and an MCMC chain length of 1 × 10^5^ with a burn-in of 1 × 10^4^. The resulting marginal likelihood estimates were used to rank the models, after which we calculated Bayes factor (BF) values, taking each of the four models as in model 1, and using the equation BF=2[MLE (model1) – MLE (model2)] (Kass & Raftery, 1995). A positive BF value suggests support for model 1 whereas a negative value shows support for model 2.

#### Ancestral range reconstruction analyses

The output of the Maximum Clade Credibility tree of the RAD-seq data run in BEAST was used to reconstruct the ancestral distribution and infer the biogeographical events in seven geographical areas (Amami, Okinawa, Miyako, Yaeyama, Taiwan, Philippines, and Australia, Fig. 1) in ingroup populations. We scored areas for monophyletic populations as follows: D (Okinawa1), E (Amami), F (Australia), G (Taiwan, Philippines, Okinawa2, Miyako, Yaeyama). Each area was treated as a possible source of the ancestral population by pre-assumption, and ABC each represents three outgroup species, *A. fissifrons*, *A. lanyuensis*, and *A. miniaceus*. We conducted ancestral range model selection to identify the best statistics for reconstruction, considering possible models (DIVALIKE, DEC, and BAYAREALIKE) implemented in BioGeoBEARs, with a *j*-parameter representing the long-distance dispersal scenario (Matzke, 2014). The biographical reconstruction analyses were implemented in Reconstruct Ancestral State in Phylogenies (RASP) v.4.2 (Yu, Blair, & He, 2020).

## RESULTS

### CO1 and RAD sequence data

We obtained 77 CO1 sequences (801 base pairs) out of 107 samples and integrated five published sequences (Su & Smith, 2014) as an outgroup. For the RAD-seq data, there were 106 samples with three outgroups (*A. lanyuensis*, *A. fissifrons*, and *A. miniaceus*). We assembled two data matrices: a 1) single-locus data set (SL_SNPs data), which is restricted to cover the first SNP per locus of the ingroup only and used for population genetic analyses; and a 2) multi-locus data set (ML_SNPs data,) which encompasses all available SNPs in each locus and which was used for phylogenetic analyses. We filtered both data sets to include 50% coverage, or occupancy rate. For SL_SNPs data, there were 107,683 loci and we kept 368 after filtering; and for ML_SNPs data, there were 468115 loci and we kept 14695 loci in the final assembly.

### Population structure and haplotype network

We tested for phenotypic differentiation, initially grouping populations per island. The results showed no variation as manifested in the scattering of samples in the PCA plot (Appendix S3-S4). A distinction of two groups, which we refer to as Amami-Okinawa Lineage (AOL) (Fig. 1b-c) and Austral-Asia Lineage (AAL) (Fig. 1b–c), mirrored results observed in STRUCTURE and CO1 TCS network analyses. Population genetic structure analyses of the SL_SNPs data revealed two differentiated lineages that correspond with the best K=2 (Fig. 1c). We also observed that *A. bonadea* in AOL is hosted by *Cyrtophora* and *Nephila* whereas *A. bonadea* in AAL exhibits a more generalist behavior, involving a variety of different hosts. To rule out the possible sample size bias, we used a rarefaction curve (Appendix S7). Our sample size for AOL lineage is 14 and its expected host species number is about 7, which is in the plateau region, indicating saturation of species number. However, the observed host species number is substantially lower, at n=2. We interpret this result as evidence for AOL kleptoparasites using only very specific host species. This indicates a significant influence of an ecological factor (i.e., host-specificity), on the genetic differentiation of these kleptoparasites. The CO1 TCS network (Fig. 1b) showed a disconnected cluster between AOL and AAL—with AOL possessing a higher haplotype diversity (0.857) than AAL (0.124) (Table 1). Our AOL cluster consists of samples from Amami and nine Okinawa samples, while AAL includes six samples from Okinawa, samples from Miyako, Yaeyama, Taiwan, Philippines and, even across Wallace’s Line, from Australia.

**Table 1:** Haplotype Diversity of two lineages.

| CO1 Cluster | Sample Size | h | Hd |
| --- | --- | --- | --- |
| AOL (Cluster 1) | 14 | 8 | 0.857 |
| AAL (Cluster 2) | 63 | 5 | 0.124 |
| Total | 77 |  |  |
| h: Number of haplotypes |  |  |  |
| Hd: Haplotype diversity |  |  |  |

### DAPC and VAE clustering results and phylogenetic trees

K-mean clustering in DAPC recovered three clusters: AOL was split into two groups, while AAL clustered as one (Fig. 2a). Results from our unsupervised machine learning analysis (VAE) supported the AOL and AAL (Fig. 2b) similar to results form STRUCTURE and TCS. Moreover, the BEAST MCC tree of ML_SNPs data (Fig. 2c) showed two strongly supported (Posterior Probability, or PP=1.00) major clades, which correspond to the two previously characterized lineages of *A. bonadea.* All Amami samples (n=7) and most of the Okinawa samples (7/11) were shown to group as one lineage (AOL), while all the other samples (four Okinawa samples, Miyako group, Yaeyama, Taiwan, Philippines, and Australia) formed a distinct large lineage (AAL) that was subdivided further into smaller groups (with lower nodal support: posterior probability < 0.5), which did not discernibly corresponded to geography. This topology is well-supported in the maximum likelihood tree (IQ-TREE 2) of ML_SNPs data (Appendix S5), with bootstrap support of 100, gCF value of 100, and a sCF value of 88; and we show the BEAST MCC tree of the CO1 genes in Appendix S6.

**Figure 2.**
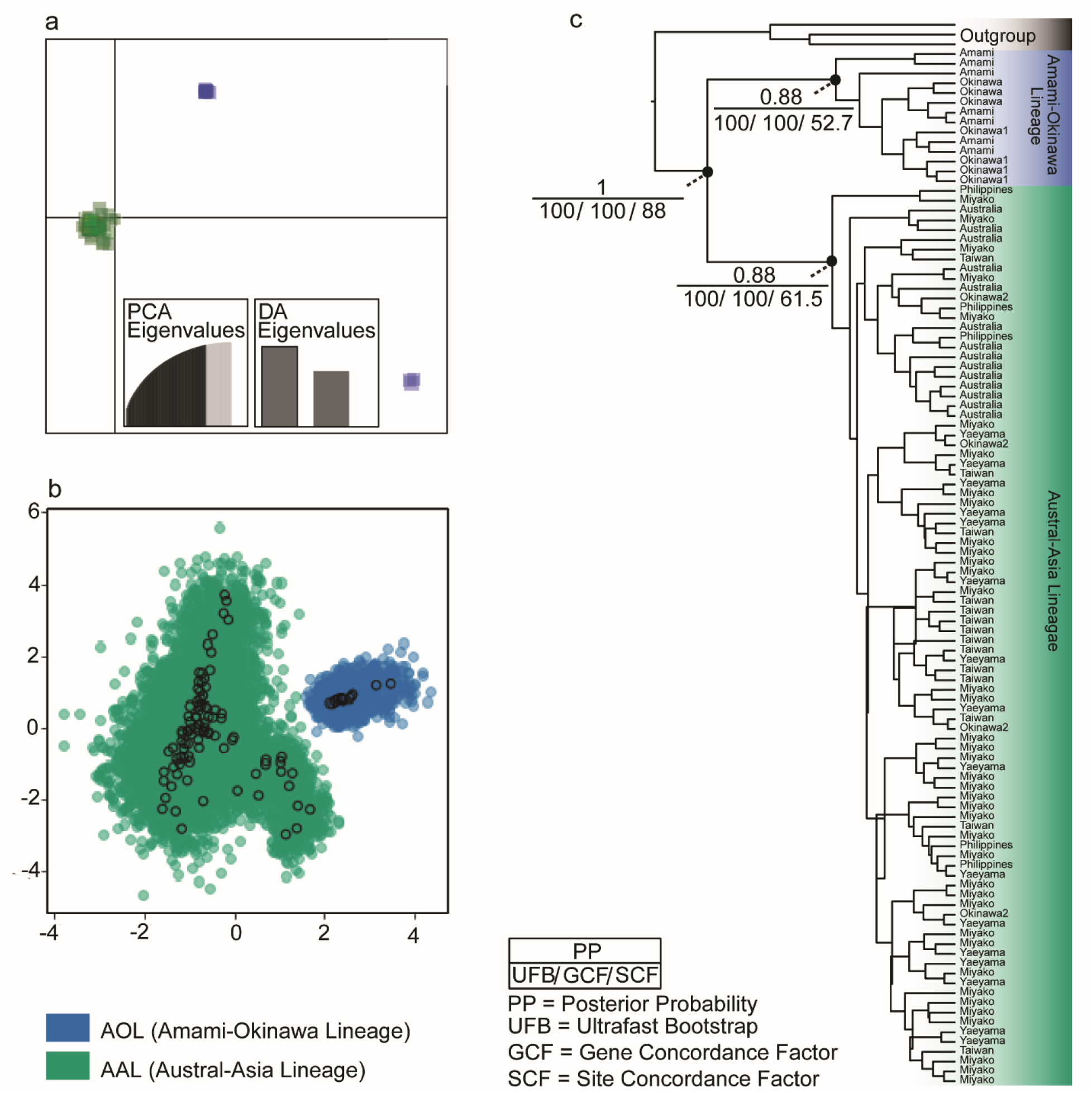
(a) K–mean clustering results using the Discriminant Analysis of Principal Components (DAPC) wherein the AOL (blue) was split into two groups (with three samples as outliers). (b) Variation Autoencoder (VAE) distinction plot between AOL and AAL (green) with an encoded mean (μ) and standard deviation (σ). (c) Maximum Clade Credibility tree of the RAD–seq data with high bootstrap support, Gene Concordance Factors (gCF), and Site Concordance Factors (sCF) at major nodes indicated (see key).

### Time calibration, species delimitation, and ancestral range reconstruction

The results of divergence time estimation analyses of SNPs data showed that the divergence of AOL and AAL occurred within the Pliocene epoch during 4.12 MYA with 0.56– 12.23 probability density range (calibrated by the divergence age of the ingroup and outgroup, Fig. 3). However, the CO1 tree (calibrated using the uncorrelated lognormal relaxed molecular clock) produced a divergence time between AOL and AAL of 25.60 MYA ago, during the early Paleocene period in the Oligocene epoch (Appendix S6), which may be unreliable, due to saturation of variation in this locus. Bayes Factor (BF) evaluation of our three hypothesized species models indicated the superiority of the one species model (Model 1 BF=-13723.4), followed by the two species model (Model 2 BF=-9956.65, Table 2). Based on these results derived from our genomic data, we consider AOL and AAL to be two lineages of the same species, which were separated via Kerama gap around 4.12 MYA.

**Figure 3.**
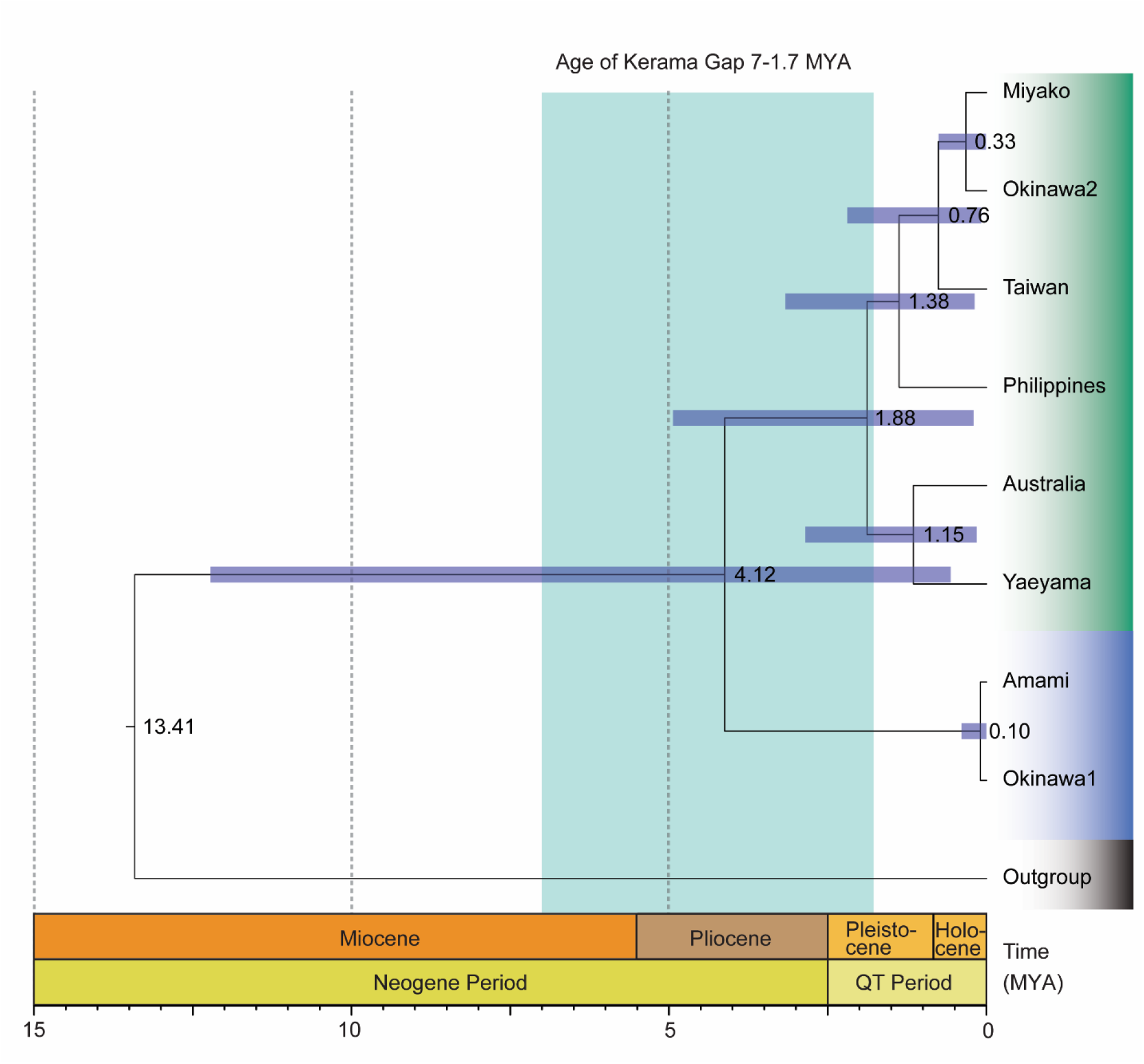
Divergence time estimation of single nucleotide polymorphisms (SNPs) data using SNAPP and calibrated based on the divergence of the outgroup and ingroup (16.18–24.02 MYA; mean 20.1 MYA; Su and Smith, 2014), and resulting in an inference of 4.12 MYA as the estimated divergence time between the Amami–Okinawa Lineage (AOL) and Austral–Asia Lineage (AAL).

**Table 2.**
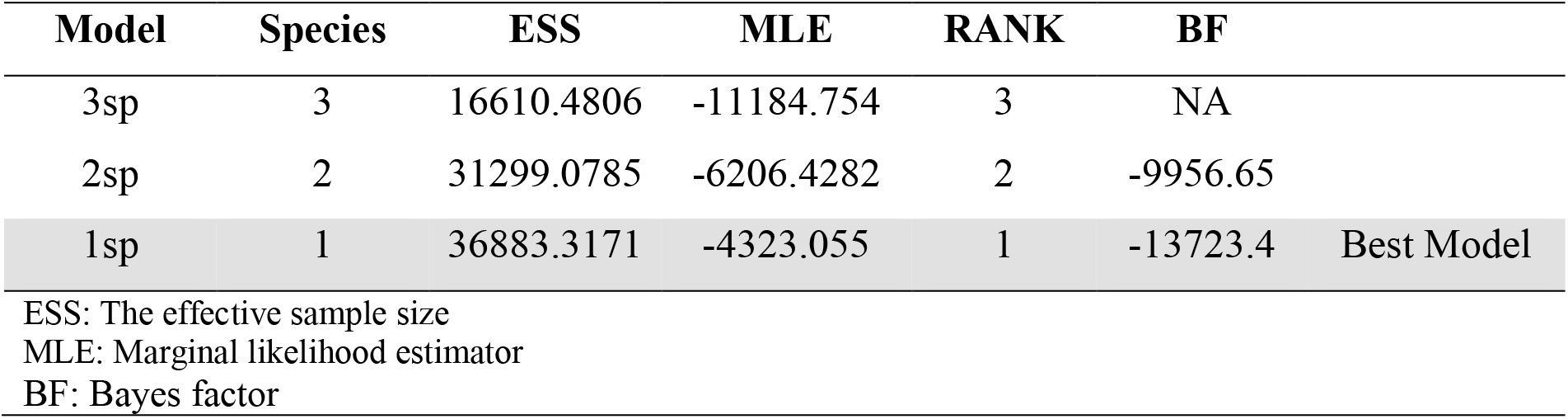
Bayes Factor Delimitation (BFD) Results.

To estimate the ancestral population and elucidate possible colonization patterns from dispersal and/or vicariance during the diversification process in *A*. *bonadea*, we considered quantitative ancestral range estimation analyses. Our RASP reconstruction of ancestral geographic range using the BEAST MCC tree identifies the DIVALIKE model as the best–fit model (AICc_wt = 0.29). The long-distance dispersal *J*-parameter is insignificant (*J* = 0). Based on the reconstructed ancestral range of node 16 (marginal probability = 31.64%; Dispersal = 1; Vicariance = 1), the ancestral range of *A*. *bonadea* most likely originates from combined areas of Taiwan, Philippines, Okinawa, Miyako and Yaeyama (DG: 37.03%). Subsequently, the ancestor at DG dispersed into the area DFG, then eventually split into the area FG or D (Event Route: DG->DFG->FG|D). A dispersal event was observed at the node splitting AOL and AAL across the Kerama Gap, an event which we estimated to be in the early Pliocene, followed by vicariance events that occurred in most of the Pliocene and until the Pleistocene, when dispersal events increased in frequency (Fig. 4).

**Figure 4.**
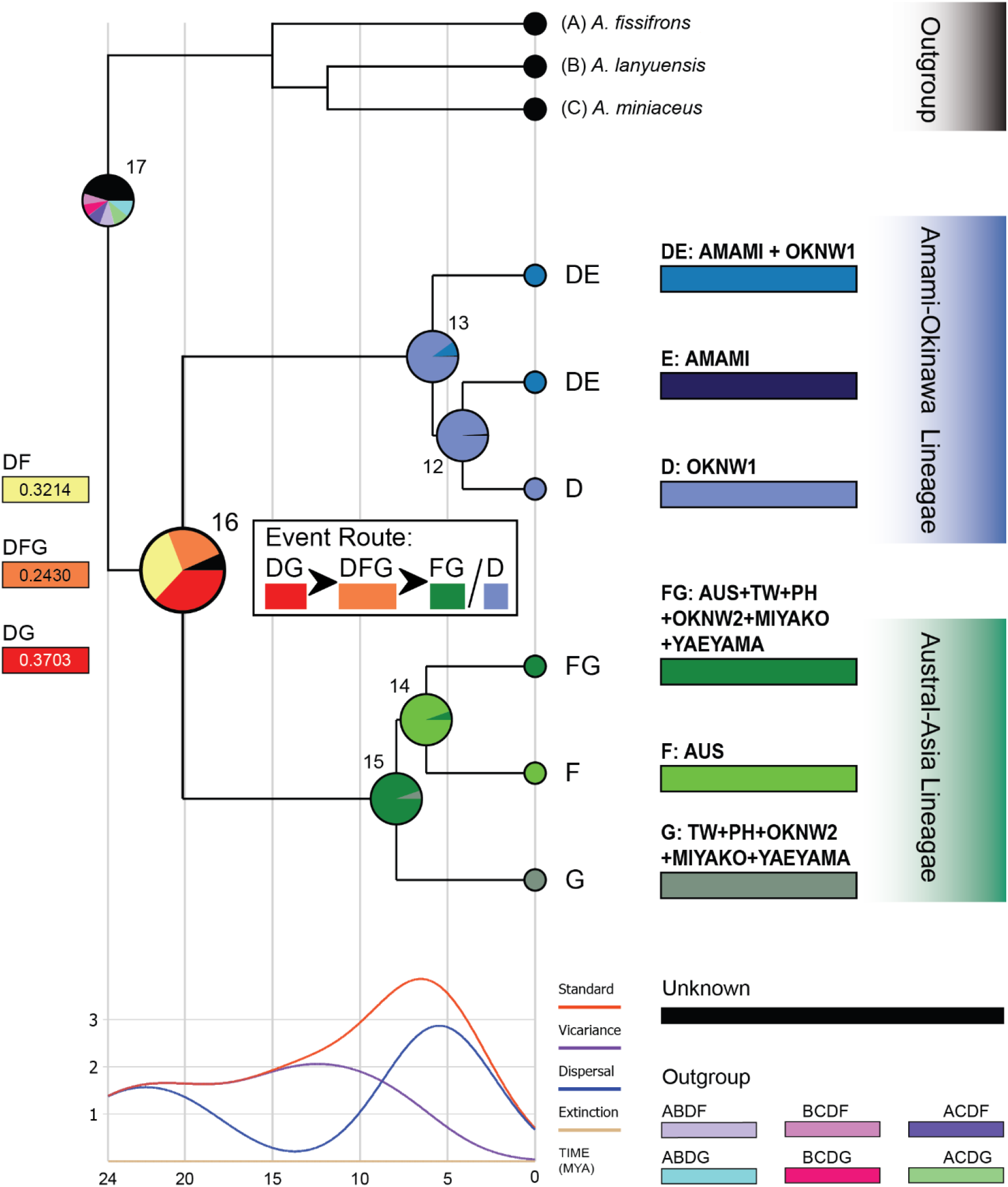
Ancestral geographic range estimates from Reconstruct Ancestral State in Phylogenies (RASP) derived from single nucleotide polymorphism (SNPs) data and a BEAST maximum clade credibility tree. The DIVALIKE model was the best–fit model; dispersal and vicariance events indicated.

## Discussion

Genetic differentiation, facilitated by long-distance dispersal, then reinforced by host adaptation, best explains the population boundary of *Argyrodes bonadea* in Taiwan-Ryukyu Islands, the Philippines, and Australia. Recent studies have thoroughly examined population diversification, speciation, and island colonization across East Asian fauna, focusing on organisms that are native to Taiwan and the Ryukyu Island Chain. (lizards: Yang et al., 2018; cicadas: Nagata, Toda, Ohbayashi, Hayashi, & Sota, 2021; cycads: Chang et al., 2022). These studies have incorporated or inferred long-distance dispersal and island colonization within variable temporal estimations. However, among such studies, choice of focal taxa confined to Taiwan and Ryukyu may have limited possible inference of other potentially significant contributing factors such as behavioral traits, or other attributes of organismal natural history. We demonstrate such a case, using a geogenomic approach, to test historical and ecological biogeographic predictions, derived from an explanatory hypothesis, incorporating both dispersal and variation in host specialization, and incorporating a broad sampling area, spanning the western Pacific, from the Ryukyu Island Chain to Australia (Fig. 1a). Our results characterize two probable incipient lineages (AOL and AAL) of *A. bonadea* from seven island populations inferred via multiple population structure and phylogenomic analyses (Fig. 1 and 2). We initially predicted that isolation of island populations should effectively hinder gene flow and we anticipated four differentiated clusters: 1) Central Ryukyu, 2) South Ryukyu and Taiwan, 3) the Philippines, and 4) Australia. However, population genetic analyses demonstrated that our two inferred lineages (AOL and AAL), roughly separated by the Karema Gap (Fig. 1a), exhibit signatures of limited between-lineage gene flow (Appendix S8); thus, their population genetic structure cannot be attributed solely to boundaries among island banks. Instead, host-specificity better explains our inference of an effective population boundary (Fig. 1c). Host species diversity appears to reduce in Amami, with *Cyrtophora* as the only observed host, whereas host specialization is more significant in Okinawa as the only island where two lineages of *A. bonadea* seems to be in sympatry. On the other hand, our inferred Austral-Asia Lineage exhibits a generalist behavior and appears tolerant of a wide range of hosts on islands of the southern Ryukyu Island Chain, Taiwan, the Philippines, and Australia (Fig. 1c). Our divergence time estimate suggests isolation of AOL and AAL during the early Pliocene (4.12 MYA, Fig. 3), likely triggered by an eastward dispersal event (Fig. 4). Our results suggest host species web specialization (a rarely-invoked category in species interactions and ecological adaptation), subsequent to long distance dispersal and island colonization, may facilitate population diversification in the kleptoparasitic spider *A. bonadea*.

Because of the documented pitfalls associated with interpretation of population structure from single-locus mitochondrial data (Zhang & Hewitt, 2003; Dong et al., 2023), the overall genetic dichotomy of AOL and AAL lineages, that was initially detected in our haplotype network analyses of CO1 data (Fig. 1b) required verification on a genomic scale. Accordingly, analyses of variation across the genome (characterization of SNPs data, using STRUCTURE and DAPC) revealed two differentiated genetic cohorts among island populations corresponding to AOL and AAL lineages, which was further reinforced by our VAE result. The strength and reliability of VAE is based on its intrinsic property being unsupervised, meaning that SNP data were input into the network system with no a priori group designation, and during the training phase, during which time the system learns patterns in the samples, then outputs clusters based on feature similarity. Overall, our clustering analyses showed clear structure, favoring two divergent genetic clusters, which contradicts our initial prediction of four clusters, confined by island group boundaries. However, when we further statistically tested the “species” status of AOL and AAL, our BFD results did not support the hypothesized two-species arrangement but, instead, supported the interpretation of one species (with a BF= −13723.4; followed by two species model [AOL and AAL], with a BF= −9956.65; Table 2), comprised of two subspecific lineages. This interpretation is also corroborated by the absence of morphological differences in the traits we measured (Appendix S3–S4) and the appearance of their genitalia (appendix S8). We calibrated our divergence time estimation based on the divergence of the outgroup and ingroup (16.18– 24.02 MYA; mean 20.10 MYA; Su and Smith, 2014) and estimated divergence between AOL and AAL at approximately 4.12 MYA (Fig. 3) when Central Ryukyu was already separated from Southern Ryukyu (Kimura, 2002). This interpretation suggests that successful colonization was accomplished even in the presence of the Kerama Gap, thus supporting overwater dispersal as a viable process underlying our inference of a structured geographical pattern (Figs. 1–3). From our time-calibrated inference of historical dispersal across the Kerama Gap, we conclude that this often-cited geographical barrier has not been an effective barrier, sufficient to prevent gene flow in *A. bonadea*. If it were, we would expect a divergence time predating 7.0 MYA, consistent with an interpretation of vicariance. It is also worth noting that the AAL shows a lack of genetic differentiation over some of the very deep channels of Sulawesi and associated islands, further suggesting that deep channels are not an effective barrier to dispersal and range expansion in this organism. Similarly, our RASP results inferred an ancestral range of AOL and AAL as Taiwan, Philippines, Okinawa, Miyako and Yaeyama (DG: 37.03%); accordingly, dispersal is supported as the likely predominant process responsible for range expansion, involving faunal exchange from the ancestral range of *A. bonadea* to distant island populations, after which subsequent differentiation may have occurred as a function of geographical isolation (Mayr,1963; Almieda et al., 2022).

Several recent studies, conducted with either genomic-scale data (Okuyama et al., 2020; Iguchi, Tada, Nagano, & Yasuda, 2021; Chang et al., 2022) or involving limited numbers of Sanger sequence loci (Su et al., 2016; Kumekawa et al., 2021) have found similar patterns, but have been restricted in geographical focus on Taiwan and the Ryukyu Island Chain. Here, we used both genome-wide, and Sanger DNA sequence data, to further extended inference of patterns to the broader regional scales of the tropical Southwest Pacific, including the Philippines and Australia (Brown et al., 2013; Lohman et al., 2011). Such a larger, more expansive geographic scope allows for the identification of additional, potentially contributing factors underlying patterns of differentiation, including organismal response to the geographical template, such as dispersal across permanent ocean barriers associated with zoogeographical regional boundaries or faunal turnover zones (e.g., Wallace Line; Evans et al., 2003; Brown, 2016; Turk, Čandek, Kralj-Fišer, & Kuntner, 2020) and the contribution of isolated refugia in generating and maintaining genetic divergence (Su et al., 2007)—much of which cannot be elucidated by the study of genetic variation at finer geographical scales.

Host-specificity, apparently an adaptative limitation of non-web building kleptoparasitic spiders coinhabiting the webs of single web-constructing spider species, is strongly suggested in this study as a potential, significantly contributing factor, which may have reinforced a genomic population boundary between two divergent lineages (AOL and AAL) in *A. bonadea*. The possible sample size bias, potentially related to a larger number of spiders sampled in the AAL, was ruled out through rarefaction curve analyses, from which our results supported the conclusion of host-specificity in the AOL. Species of the genus *Argyrodes* are generally categorized, based on host species webs they inhabit, as either generalists (parasitizing multiple web-building spider species) or specialists (limited to a single web-builing spider species’ webs; Vollrath, 1979). Generalists usually thrive on different kinds of hosts, whereas specialists are found on a single species’ web, or a few types of hosts (Whitehouse, 1988). *Argyrodes bonadea* is a generalist kleptoparasite but our novel inference of the AOL lineage in this study demonstrates a “specialist” type of host preference, in which the Amami population thrives exclusively on the webs of species of the spider genus *Cyrtophora* but Okinawa samples are hosted solely on webs of members of the genus *Nephila*. Together, we hypothesize that these fine scale host preferences may have led to genetic differentiation in these isolated island populations. In contrast, allopatric island populations of the broadly distributed AAL elucidated here are found on a wide variety of different host species’ webs, and across a broad expanse of the southwest Pacific. Together, these populations demonstrate tolerance of multiple host species’ webs, in the rapidly-range-expanded AAL, which may facilitate or contribute to gene flow, as manifested by minimal genetic differentiation, similar to a previously documented case, in the related island endemic kleptoparasite, *Argyrodes lanyuensis* (Responte, et al., 2021).

## CONCLUSION

In this study, we employed a geogenomics approach to elucidate the role of long-distance dispersal (as opposed to vicariance) and host adaptation in shaping genetic structure of the highly dispersive kleptoparasitic spider *A. bonadea*. Populations of this unusual spider overcame deep-sea barriers and remarkable overseas distances, giving rise to a pattern of diversification which stretches approximately a thousand kilometers, from the Ryukyu Island Chain to Australia. In a geographical pattern of genetic variation similar to that observed in one of its hosts (*Nephila pilipes*; Su et al., 2007), we found no pronounced genetic structure, or association of genetic variation with geographic distance, even at the unprecedented scale of the entire southwestern Pacific. Surprisingly, at much more limited geographically local scales of the central Ryukyu archipelago, we found substantial genetic differentiation in association with—and interpreted here as likely to have resulted from host-specificity. Although this novel empirical finding provides initial evidence for what we assume will be an enhanced appreciation of the significant influence of host transitions and/or host-specificity on population genetic structure in kleptoparasites, identification of regions of the genome or specific loci responsible for population level genetic variation, will be an important future priority, necessary to gain richer, more nuanced understanding of both the proximate cause and ultimate adaptive significance of this variation.

## ACKNOWLEDGMENTS

In gratitude, we acknowledge the members of the Ecology and Evolutionary Genomics (EEG) laboratory at Kaohsiung Medical University for their assistance in the development and completion of this work. We thank the Biodiversity Management Bureau (BMB; formerly Protected Areas and Wildlife Bureau, PAWB) of the Department of the Environment and Natural Resources (DENR) in the Philippines for allowing YCS to collect and export samples under the aegis of a Gratuitous Permit (No. 171) to Collect Biological Specimens and an export permit issued to (RMB; supported by NSF DEB 0743491). The samples collected in Taiwan were under the permit (to YCS) of the Agriculture Department of Taitung County Government (June 2019). Australian samples were collected under the permit (to PP) of Brisbane Infrastructure Field Services, Asset Services North (December 2018 to January 2019). We are grateful to Ting-Siou Chen and Wan-Ching Huang for the fieldwork in Ryukyu Island Chain, Japan in February 2020. We received valuable information and support in complying with Japanese laws and regulations on biodiversity conservation, from the ABS Support Team at Makino Herbarium, Tokyo Metropolitan University (Leader: Noriaki Murakami in FY2017–2021; Katsuyuki Eguchi in FY2022–2026) under the National BioResource Project of the Ministry of Education, Culture, Sports, Science and Technology (MEXT), Japan. We are also thankful to Charlyn Villavicencio-Rosales for helping us with computer command troubleshooting. The animal handling protocols were approved by Kansas University institutional animal care and use committee (IACUC auth. 185-05 to RMB). This work was supported by the National Science and Technology Council (NSTC), Taiwan (funding number 107-2311-B-037-004-MY3 and 110-2621-B-037-001-MY3).

## DATA AVAILABILITY

The data used to support the findings of in this study are openly available in Dryad at https://doi.org/10.5061/dryad.1c59zw40r

The new DNA sequences can be accessed from GenBank (OQ748070).

## BIOSKETCH

**Noraya U. Elias** is a faculty member of the Mindanao State University-Malabang Community High School, Malabang, Lanao del Sur, Philippines and currently a Ph.D. student at the Ecology and Evolutionary Genomics (EEG) laboratory at the Department of Biomedical Sciences and Environmental Biology, College of Life Science, Kaohsiung Medical University, Kaohsiung, Taiwan. Her Ph.D. research focuses on kleptoparasitic spider, *Argyrodes bonadea* as a model to test the importance of the historical biogeographic processes and the ecological factors in the differentiation/ speciation process. The research team in EEG laboratory headed by **Dr.Yong-Chao Su** utilizes genomic tools to answer questions in ecological adaptation, phylogenetics, biogeography, and functional trait evolution of which insects and spiders are often used as research materials.

