## Supplemental File 1 for "A shift in the host web occupancy of dew-drop spiders associated with genetic divergence in the Southwest Pacific": Supplementary Materials.docx

**Appendices**

Appendix S1. *Argyrodes bonadea* and host web-species association data. See text for methods of host species determination and other methodological details of characterizing natural history traits in the field (Su et al*.*, 2014).

| Sample ID | Host | Location |
| --- | --- | --- |
| EEG 783 | *Cyrtophora* | Amami |
| EEG 784 | *Cyrtophora* | Amami |
| EEG 785 | *Cyrtophora* | Amami |
| EEG 941 | *Cyrtophora* | Amami |
| EEG 942 | *Cyrtophora* | Amami |
| EEG 943 | *Cyrtophora* | Amami |
| EEG 944 | *Cyrtophora* | Amami |
| EEG 710 | *Nephila* | Okinawa |
| EEG 711 | *Nephila* | Okinawa |
| EEG 712 | *Nephila* | Okinawa |
| EEG 719 | *Nephila* | Okinawa |
| EEG 720 | *Nephila* | Okinawa |
| EEG 777 | *Nephila* | Okinawa |
| EEG 714 | *Cyclosa* | Okinawa |
| EEG 716 | *Cyclosa* | Okinawa |
| EEG 717 | *Cyclosa* | Okinawa |
| EEG 718 | *Cyclosa* | Okinawa |
| EEG 924 | *Cyrtophora* | Yaeyama |
| EEG 925 | *Cyrtophora* | Yaeyama |
| EEG 926 | *Cyrtophora* | Yaeyama |
| EEG 927 | *Cyrtophora* | Yaeyama |
| EEG 928 | *Cyrtophora* | Yaeyama |
| EEG 929 | *Cyrtophora* | Yaeyama |
| EEG 930 | *Cyrtophora* | Yaeyama |
| EEG 932 | *Cyrtophora* | Yaeyama |
| EEG 933 | *Cyrtophora* | Yaeyama |
| EEG 934 | *Cyrtophora* | Yaeyama |
| EEG 935 | *Cyrtophora* | Yaeyama |
| EEG 936 | *Cyrtophora* | Yaeyama |
| EEG 937 | *Cyrtophora* | Yaeyama |
| EEG 938 | *Cyrtophora* | Yaeyama |
| EEG 939 | *Cyrtophora* | Yaeyama |
| EEG 940 | *Cyrtophora* | Yaeyama |
| EEG 700 | *Neoscona* | Miyako |
| EEG 701 | *Nephila* | Miyako |
| EEG 702 | *Nephila* | Miyako |
| EEG 703 | *Nephila* | Miyako |
| EEG 704 | *Nephila* | Miyako |
| EEG 705 | *Nephila* | Miyako |
| EEG 706 | *Nephila* | Miyako |
| EEG 707 | *Nephila* | Miyako |
| EEG 708 | *Nephila* | Miyako |
| EEG 945 | *Nephila* | Miyako |
| EEG 946 | *Nephila* | Miyako |
| EEG 947 | *Nephila* | Miyako |
| EEG 948 | *Nephila* | Miyako |
| EEG 949 | *Nephila* | Miyako |
| EEG 950 | *Parasteatoda* | Miyako |
| EEG 799 | *Nephila* | Miyako |
| EEG 922 | *Leucauge* | Miyako |
| EEG 923 | *Nephila* | Miyako |
| EEG 800 | *Parasteatoda* | Ikema |
| EEG 854 | *Parasteatoda* | Ikema |
| EEG 855 | *Parasteatoda* | Ikema |
| EEG 856 | *Parasteatoda* | Ikema |
| EEG 857 | *Parasteatoda* | Ikema |
| EEG 858 | *Nephila* | Ikema |
| EEG 859 | *Nephila* | Ikema |
| EEG 860 | *Nephila* | Ikema |
| EEG 861 | *Nephila* | Ikema |
| EEG 862 | *Nephila* | Irabu |
| EEG 863 | *Nephila* | Irabu |
| EEG 864 | *Nephila* | Irabu |
| EEG 866 | *Nephila* | Irabu |
| EEG 913 | *Nephila* | Irabu |
| EEG 914 | *Nephila* | Irabu |
| EEG 915 | *Nephila* | Irabu |
| EEG 916 | *Nephila* | Irabu |
| EEG 917 | *Nephila* | Irabu |
| EEG 918 | *Nephila* | Irabu |
| EEG 919 | *Nephila* | Irabu |
| EEG 920 | *Nephila* | Irabu |
| EEG 921 | *Nephila* | Irabu |
| EEG 1227 | *Cyrtophora* | Taiwan |
| EEG 1229 | *Cyrtophora* | Taiwan |
| EEG 1230 | *Cyrtophora* | Taiwan |
| EEG 1231 | *Cyrtophora* | Taiwan |
| EEG 1233 | *Cyrtophora* | Taiwan |
| EEG 1234 | *Cyrtophora* | Taiwan |
| EEG 1235 | *Cyrtophora* | Taiwan |
| EEG 1236 | *Cyrtophora* | Taiwan |
| EEG 1237 | *Cyrtophora* | Taiwan |
| EEG 1238 | *Cyrtophora* | Taiwan |
| EEG 1240 | *Cyrtophora* | Taiwan |
| EEG 1264 | *Cyrtophora* | Taiwan |
| EEG 1269 | *Cyrtophora* | Taiwan |
| EEG 786 | *Nephila* | Philippines |
| EEG 787 | *Nephila* | Philippines |
| EEG 788 | *Argiope* | Philippines |
| EEG 789 | *Nephila* | Philippines |
| EEG 958 | *Nephila* | Philippines |
| EEG 790 | *Cyrtophora* | Australia |
| EEG 791 | *Cyrtophora* | Australia |
| EEG 792 | *Cyrtophora* | Australia |
| EEG 793 | *Cyrtophora* | Australia |
| EEG 794 | *Cyrtophora* | Australia |
| EEG 795 | *Cyrtophora* | Australia |
| EEG 796 | *Cyrtophora* | Australia |
| EEG 797 | *Cyrtophora* | Australia |
| EEG 952 | *Cyrtophora* | Australia |
| EEG 953 | *Cyrtophora* | Australia |
| EEG 954 | *Cyrtophora* | Australia |
| EEG 955 | *Cyrtophora* | Australia |
| EEG 956 | *Cyrtophora* | Australia |
| EEG 957 | *Cyrtophora* | Australia |

Appendix S2. Detailed methods and processing pipeline for RAD-seq library preparation and sequence assembly.

The protocol for Multiplexed Shotgun Genotyping (MSG)-Illumina Deep Sequencing Mapping was carried out (Andolfatto et al*.*, 2011) to prepare the RAD-seq libraries. Initially, 10μl of 25ng of each DNA sample was prepared and then digested in the digestion master mix containing 7.7μl nuclease-free water, 2μl of 10X cutsmart and 0.3μl of MseI restriction enzyme. Incubation was at 37°C for 3 hours followed by inactivation at 65°C for 20 minutes then ligated to a unique bar-coded adapter at 16°C for 3 hours. Then ligated samples were pooled and precipitated in 99.8% ethanol and chilled overnight at 4°C. Samples were washed repeatedly with 70% ethanol and then resuspended with 40μl of TE buffer. Size selection was done in a Pippin Prep machine (Pippin Prep, Sage Science, Inc., USA) to target a genomic fragment of 375 base pairs. Amplification of the barcoded fragments was done using the Phusion PCR kit and FC1 and FC2 primers. Purified final products were sent to Genomics Co. Ltd., Taiwan for Illumina NovaSeq 6000 sequencing with 2 x 150 bp kit.

We used Stacks v2.0 (Catchen et al*.*, 2013) to assemble the raw RAD-seq sequences. Stacks assembly operates in three major steps: (1) the process_radtags; the gstacks; and (3) the populations. In process_radtags, the genomic library is sorted, cleaned, and demultiplexed according to the pre-specified oligo-barcode. Then Paired-End read merger (Zhang, Kobert, Flouri, & Stamatakis, 2014) was used to merge the forward and reverse strands. The paired strands were aligned to the *Argyrodes miniaceus* genome draft sequenced via Hi-seq 2500 platform as the reference with the aid of a standard alignment program Burrows-Wheeler Aligner (BWA). The gstacks program processes the aligned data and calls SNPs using the population-wide data per locus as executed in ref_map.pl. The population program filters the data, calculates the population genetics statistics and produces different data formats to be used for the downstream analyses. We defined population as a set of individuals per island. Loci were filtered out at 50% occupancy rate to include locus that is present in at least half of the total individuals in a population.

Appendix S3. Bivariate ordination of the first two (of three total) principal components, extracted in multivariate analysis of 15 continuous morphological characters, measured from male specimens and demonstrating phenotypic homogeneousity and the absence of group-based structure (i.e., island population distinctiveness) in phenotypic morphospace. We measured the total length (TL), carapace length (CL), carapace width (CW), abdomen length (AL), abdomen width (AW), abdomen height (AH), total length of each leg (I-IV), and the length of each leg (I-IV) segments. The palpal length (PL) of the male samples was also measured. All the morphometric measurements were standardized according to CL. The carapace is generally conserved in shape especially among species of the same family (Foellmer & Moya-Laraño, 2007; Kallal, Moore, & Hormiga, 2019).

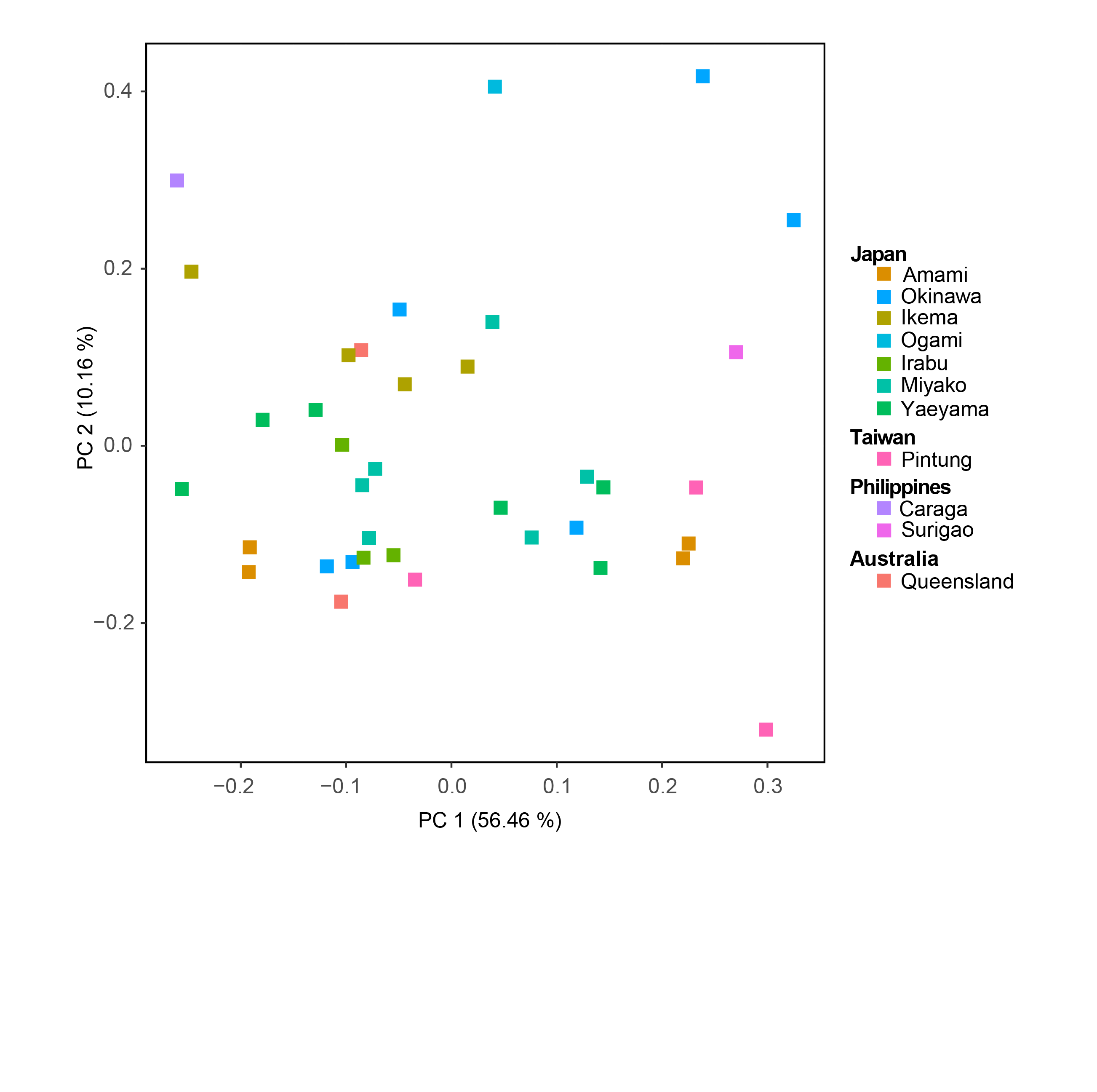

Appendix S4. PCA plot of morphometrics of female samples which shows phenotypically homogeneous. Characters and abbreviations follow those provided in Appendix S3. We calculated the total length, carapace length, carapace width (CW), abdomen length (AL), abdomen width (AW), abdomen height (AH), total length of each leg (I-IV), and the length of each leg (I-IV) segments. The palpal length (PL) of the male samples was also measured. All the morphometric measurements were standardized according to CL. The carapace is generally conserved in shape especially among species of the same family (Foellmer & Moya-Laraño, 2007; Kallal et al., 2019)

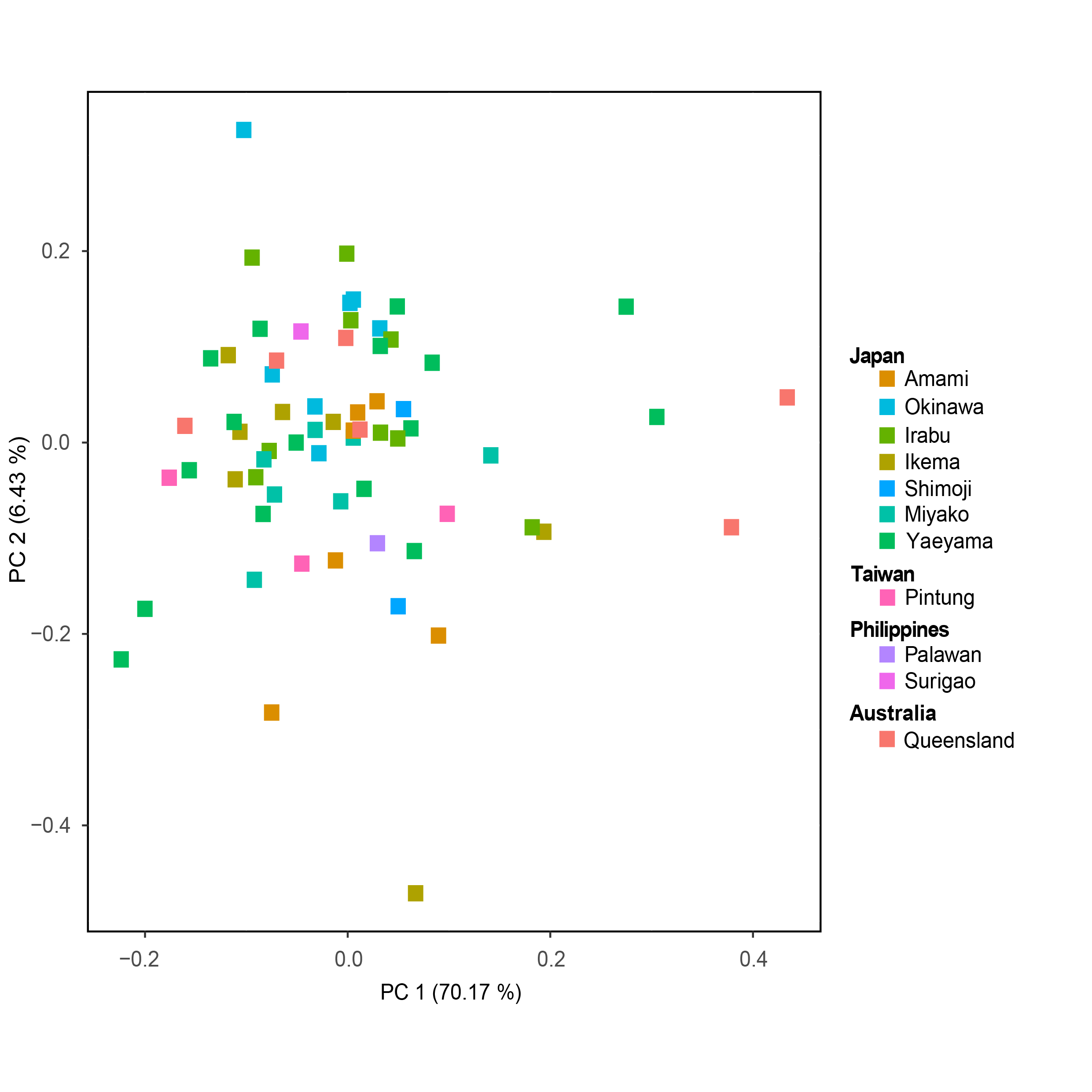

Appendix S5. Alternative maximum-likelihood reference tree constructed in IQ-TREE 2 using RADseq (ML SNPSdata) with a strong Ultrafast Bootstrap (UFB) support node between the Amami-Okinawa Lineage and Austral-Asia Lineage (>95), Gene Concordance Factor (GCF) and Site Concordance Factor (SCF) values were >50% which showed support for congeneric taxa.

Appendix S6. Bayesian consensus tree summarizing the last one million generations from the posterior distribution in our analysis of 801 base pairs of the Cytochrome Oxidase 1 mitochondrial gene region as performed in the program BEAST (Drummond et al*.*, 2012). Major clades showed strong Posterior Probability support (PP = 1). Divergence time in million years (MY) with 95% height posterior density shown as broad horizontal bars.

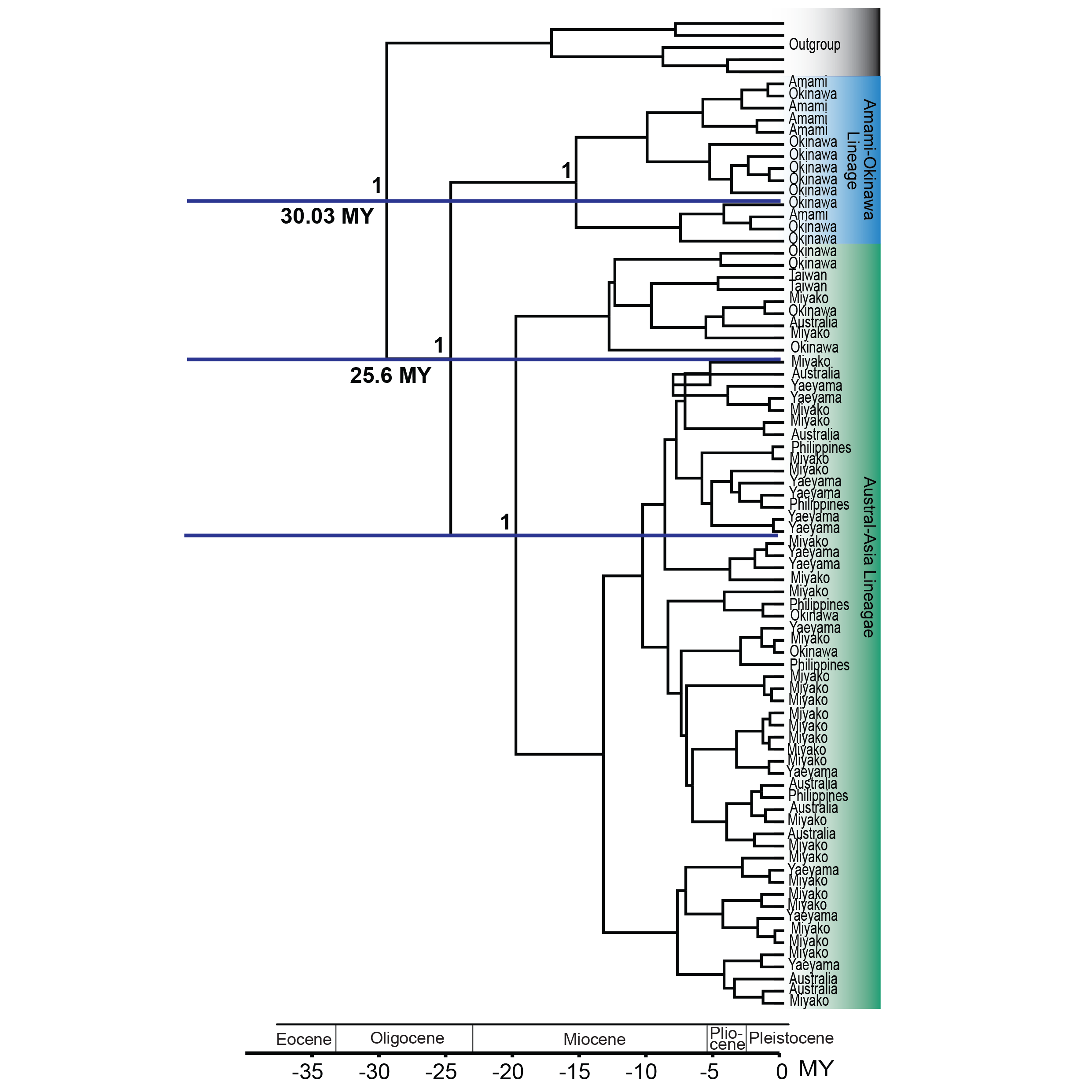

Appendix S7. Rarefaction curve shows that the expected host species number is 7 which is higher than the observed host species number which is 2, suggesting that Amami-Okinawa Lineage have strong preference for very specific host species.

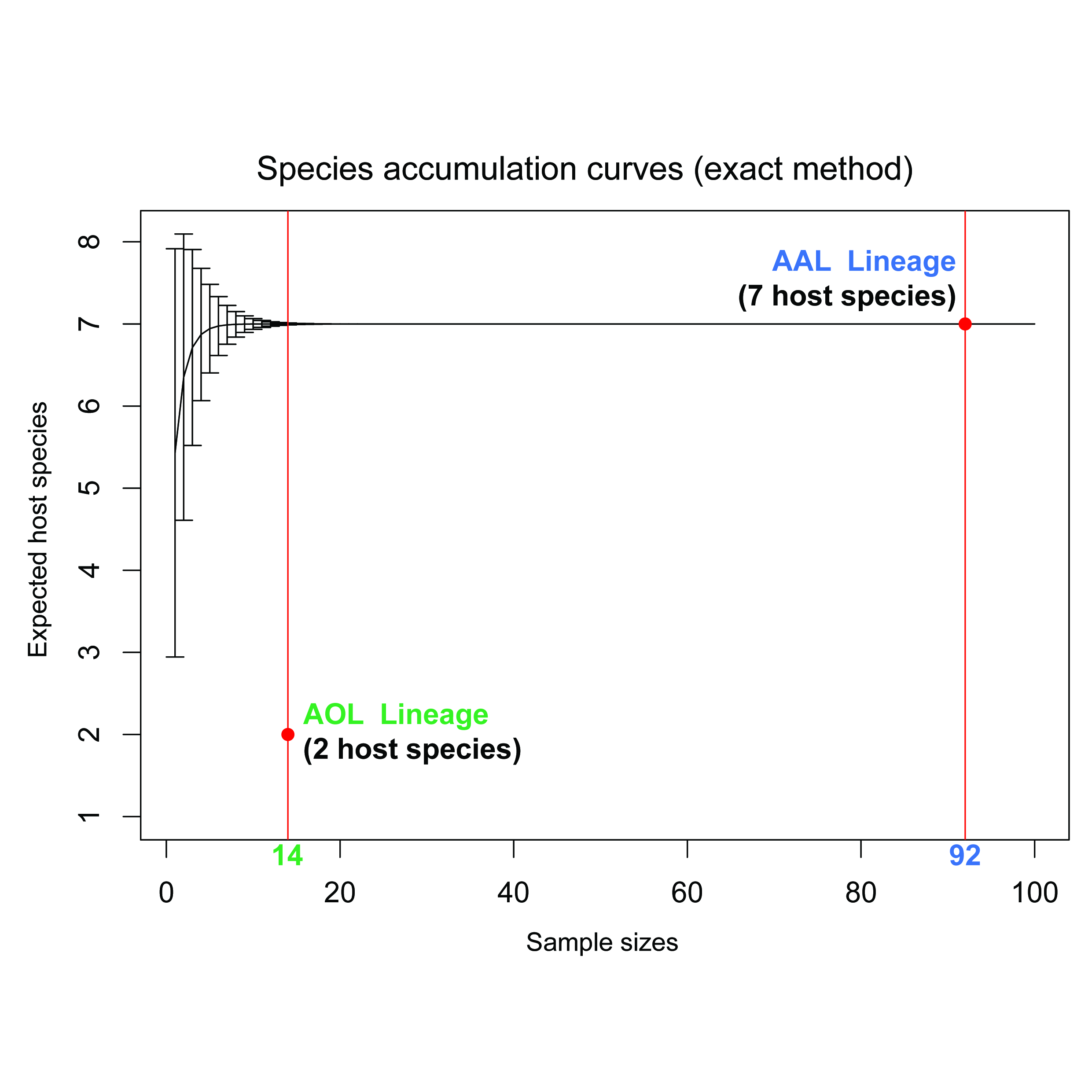

Appendix S8. Nei’s Pairwise Fst values demonstrating limited geographical structure between Amami, Okinawa and other island populations.
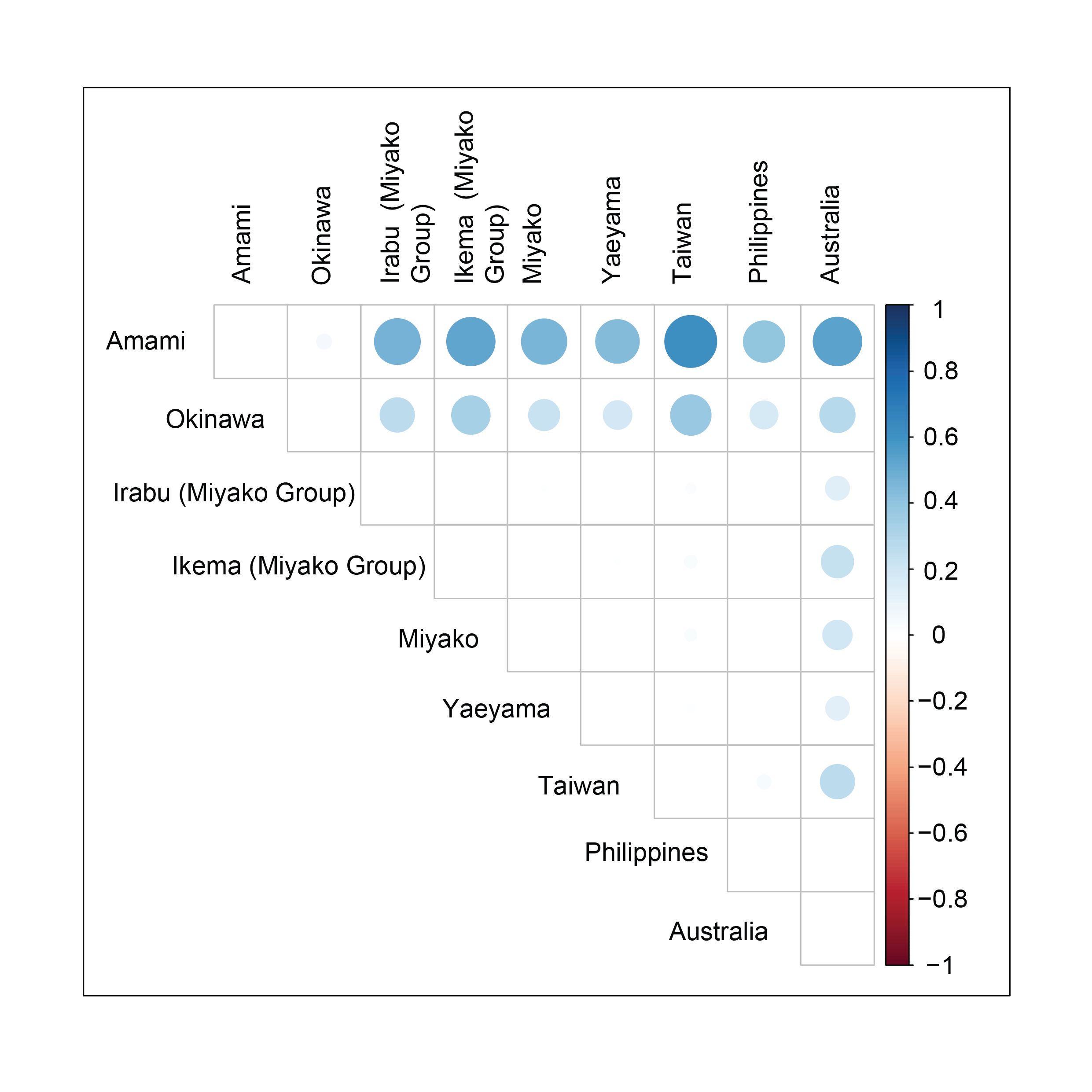

Appendix S9. Microscopic photos of the genitalia show no significant difference between the two lineages. (a) Epigynum and (b) pedipalp of the Amami-Okinawa Lineage; (c) Epigynum and (d) pedipalp of the Austral-Asia Lineage.
